# Using Relion software within Scipion framework

**DOI:** 10.1101/2020.12.06.399808

**Authors:** Grigory Sharov, Dustin R. Morado, Marta Carroni, José Miguel de la Rosa-Trevín

**Affiliations:** MRC Laboratory of Molecular Biology, Cambridge Biomedical Campus, England; Department of Biochemistry and Biophysics, Science for Life Laboratory, Stockholm University, Stockholm, Sweden

**Keywords:** Scipion, Relion, image processing, cryo-EM, SPA

## Abstract

Scipion is a modular image processing framework integrating several software packages under a unified interface while taking care of file formats and conversions. Here new developments and capabilities of the Scipion plugin for the Relion software are presented and illustrated with the image processing pipeline of published data. The user interfaces of Scipion and Relion are compared and the key differences highlighted, allowing this manuscript to be used as a guide for both new and experienced users of these software. Different streaming image processing options are also discussed demonstrating the flexibility of the Scipion framework.

**Synopsis:** An overview of the Scipion plugin for the Relion software is presented and various capabilities of image processing within Scipion framework are discussed.

## 1. Introduction

The cryogenic electron microscopy (cryo-EM) field continues to grow rapidly due to constant improvements in both instrumentation and software algorithms (Danev et al., 2019). With more high-resolution structures being solved every year (https://www.ebi.ac.uk/pdbe/emdb/statistics_main.html/) by single-particle analysis (SPA) and subtomogram averaging methods, automation of existing image processing pipelines becomes more relevant. Moreover, the introduction of user-friendly graphical interfaces (GUI) plays an important role in attracting more scientists to the field. Scipion (de la Rosa-Trevín et al., 2016) is an integrative software suite that aims to reach a larger scientific audience by providing a convenient modular platform for building custom automatable pipelines. The platform can be adapted to suit specific image processing needs of structural biologists, electron microscopy facilities or software developers. The Scipion framework includes multiple modules (plugins) each providing a set of Python wrappers around a particular cryo-EM software. Scipion collects these modules within a unified GUI allowing for seamless transition between different packages and file formats. Here we describe in detail the Scipion plugin for the Relion software (Zivanov et al., 2019) and illustrate the advantages of the image processing using the Scipion platform.

## 2. Plugin overview

With the high popularity of Relion within the cryo-EM community (https://www.ebi.ac.uk/pdbe/emdb/statistics_main.html/) the plugin for Relion is one of the most widely used by Scipion users (http://scipion.i2pc.es/report_protocols/protocolTable/). Recently the plugin has undergone several important changes. First, the code has been migrated to Python 3 since Python 2.7 was deprecated in January 2020. While being entirely transparent to users, this transition has brought many improvements for software developers and allowed us to keep the code base up-to-date. Second, a new STAR file parser *emtable* (de la Rosa-Trevin and Sharov, 2020) was developed to simplify and speed up metadata conversion between Scipion and Relion, replacing the functions of the plugin for XMIPP (de la Rosa-Trevín et al., 2013), previously used for this task. *emtable* is available as a small self-contained Python module and can be used to manipulate STAR files independently from Scipion.

Considerable effort was put into compatibility between different Relion versions. Starting with Relion version 3.1, information about optic groups has been added as a second data table to STAR files. Optic groups were designed to keep all information related to the data acquisition (pixel size, voltage etc.) separate from the particle metadata and to allow users to combine different datasets. Despite the fact that we no longer provide support for Relion 3.0 or older versions, users are still able to import older STAR files into Scipion and then continue with Relion 3.1. Most plugin protocols are also likely to work with Relion 3.0. We have created a special protocol called “assign optics groups” to account for the differences between the Scipion EM data model and new optics groups data table. Users can assign the parameters of one or more (using an extra STAR file) optics groups to a set of images at essentially any step of the image processing pipeline. This allows more flexibility compared to the Relion GUI where users have to specify optics groups at the import stage.

At the moment the plugin provides wrappers for most of the programs available from Relion (version 3.1) with the exception of protocols for helical image processing (He and Scheres, 2017) and subtomogram averaging (Bharat et al., 2015) which are still in development.

## 3. User interface design

The original idea behind a Scipion wrapper for Relion programs was to make the user interface similar to the original Relion GUI so that Relion users starting to use Scipion could easily orient themselves through many available options. Below we describe the interface of the plugin for Relion and compare it to the Relion GUI (fig. 1).

**Figure 1.**
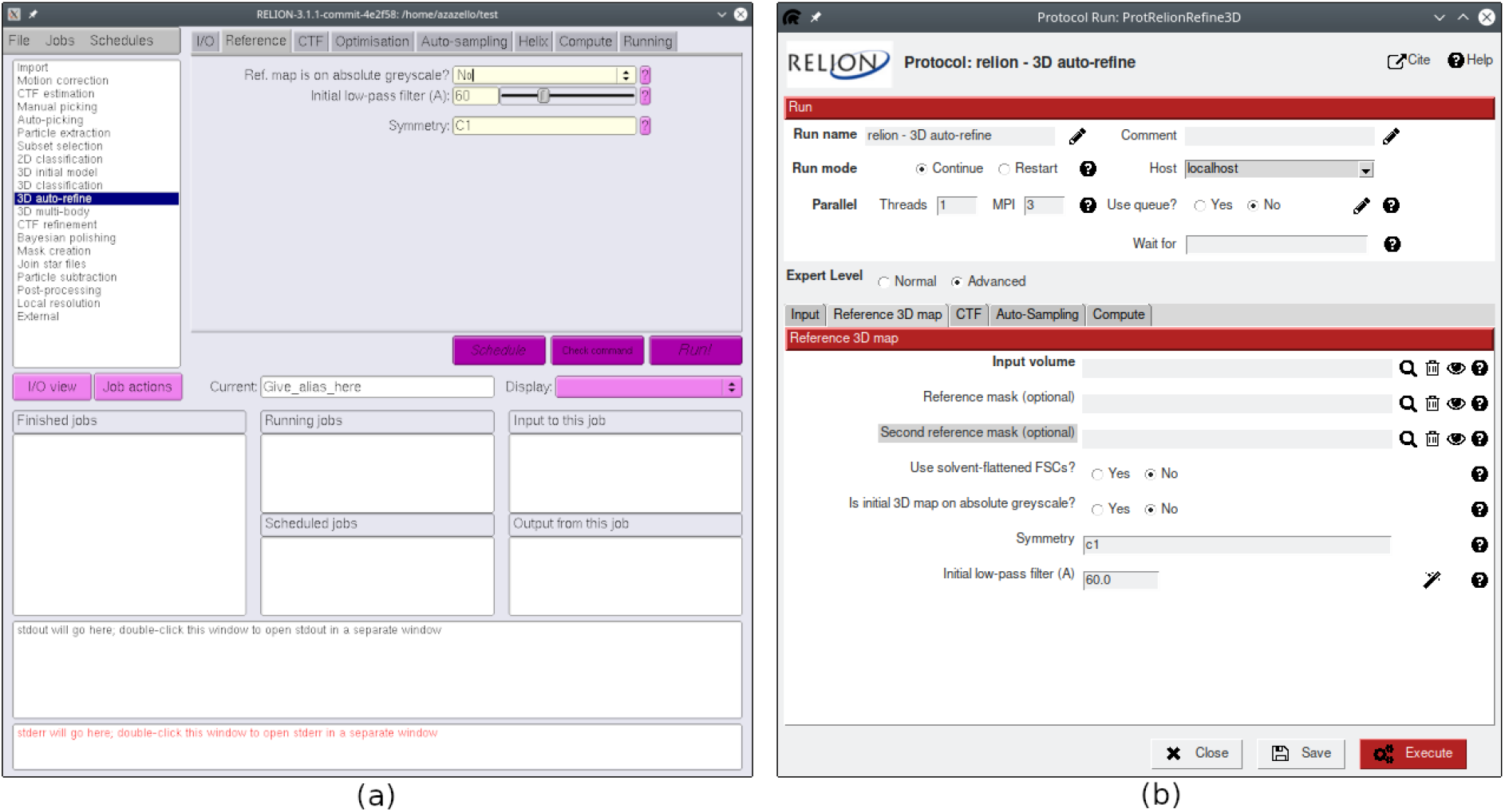
Comparison of the 3D auto-refine protocol GUI in Relion (a) and Scipion (b). Advanced protocol parameters in Scipion are highlighted with gray background.

Every protocol window in Scipion has a similar layout composed of two parts. The top panel includes computation-related parameters such as submission to a cluster queue, waiting list (the current protocol will not start until the listed protocols have finished) and parallelization options (GPU IDs, number of MPI processes and/or threads) if the protocol supports them. Furthermore, many protocols define two levels of user expertise: normal and advanced (see “Expert Level” in fig. 1). Selecting the advanced level displays additional parameters that are less common or reserved for special cases. Many plugin protocols also offer a string-type variable called “Extra parameters” or “Additional arguments”. Such strings are reserved for Relion command-line options and arguments that are not available through the standard GUI and can be utilized by advanced Relion users.

The bottom panel of the protocol window includes task-specific parameters as found in the Relion GUI, organised in similarly named tabs. Some of the parameters have an interactive wizard attached (fig. 2), meaning that one can quickly visualize how changing certain values might affect the results. Most wizards are used to select a particle mask size or to choose image filtering levels, however there are also special cases like choosing a detector MTF file or creating defocus groups.

**Figure 2.**
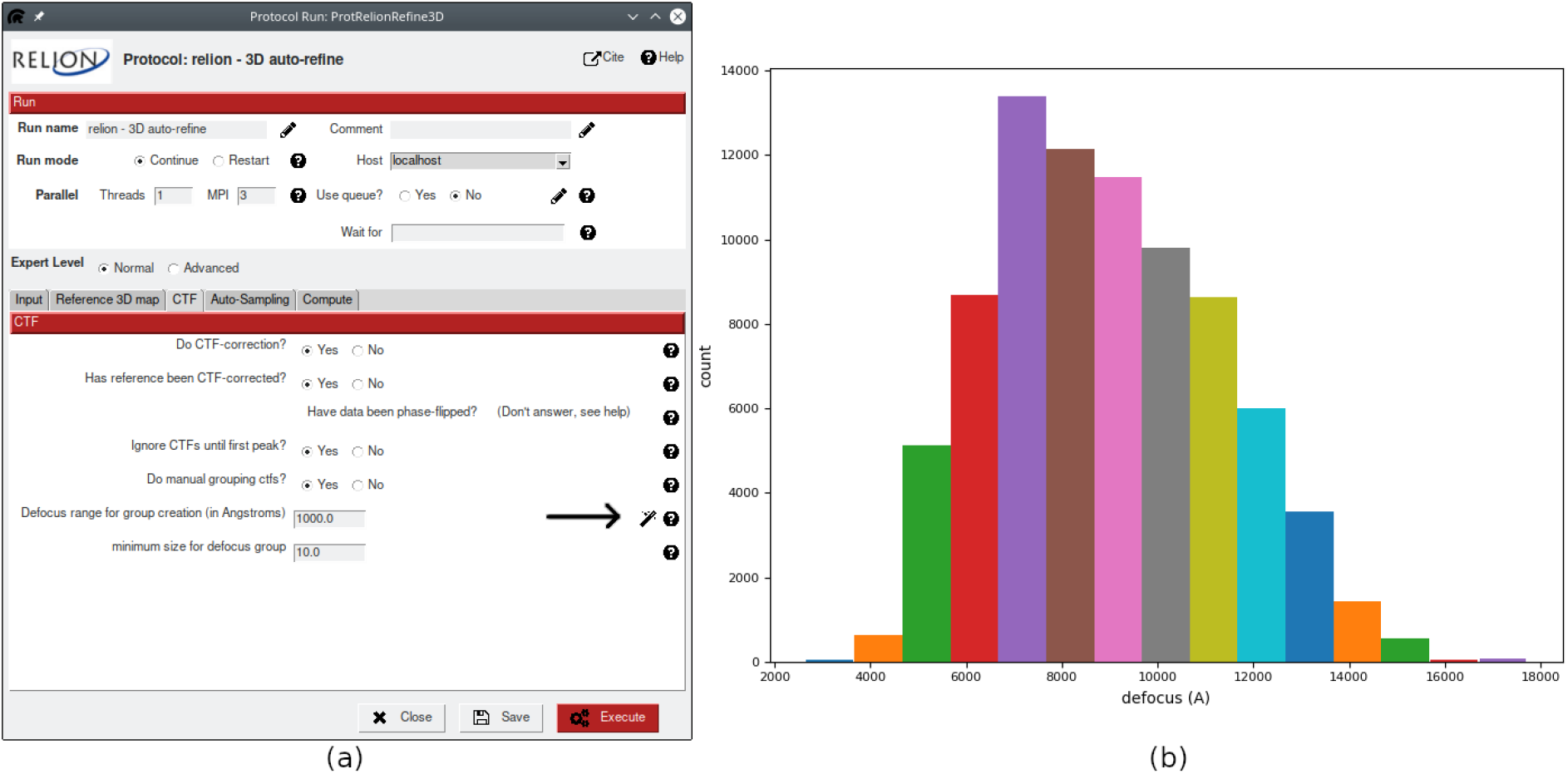
The plugin for Relion provides interactive wizards to help users choose optimal parameters. Arrow (on the (a)) indicates the wizard button that launches a histogram plot (b) with an estimated number of CTF groups given the input defocus range.

Despite many similarities with the Relion GUI, the plugin for Relion software has several differences due to a conceptually different design of the Scipion framework. One example is the “continue” mode for classification/refinement protocols. While Relion allows continuation of a finished job, in Scipion every run is unique and the user must create a new protocol which should reference the previous one. In this way the meaning of the number of iterations becomes different between Relion and Scipion: for Relion it’s a total number of iterations, but for Scipion it’s a number of extra “continue” iterations.

Another example is subset selection, which is done in Relion with a separate job type. In Scipion subsets can be created naturally from any set of items in an interactive manner. Scipion uses XMIPP viewer to display sets of images and associated metadata (fig. 3a). These sets can be displayed as an image gallery or as a table, where each column’s values can be sorted or plotted using built-in tools. This means any STAR file can be visualised with Scipion without using a text editor. Moreover, Scipion includes a set of protocols that can perform various operations on image sets such as joining, splitting or intersection. When displaying single items like a micrograph or a particle, Scipion offers a range of ImageJ (Schneider et al., 2012) image processing tools that can be used, for example, to calculate a fast Fourier transform (FFT) or to interactively create a 2D mask from an image (fig. 3b).

**Figure 3.**
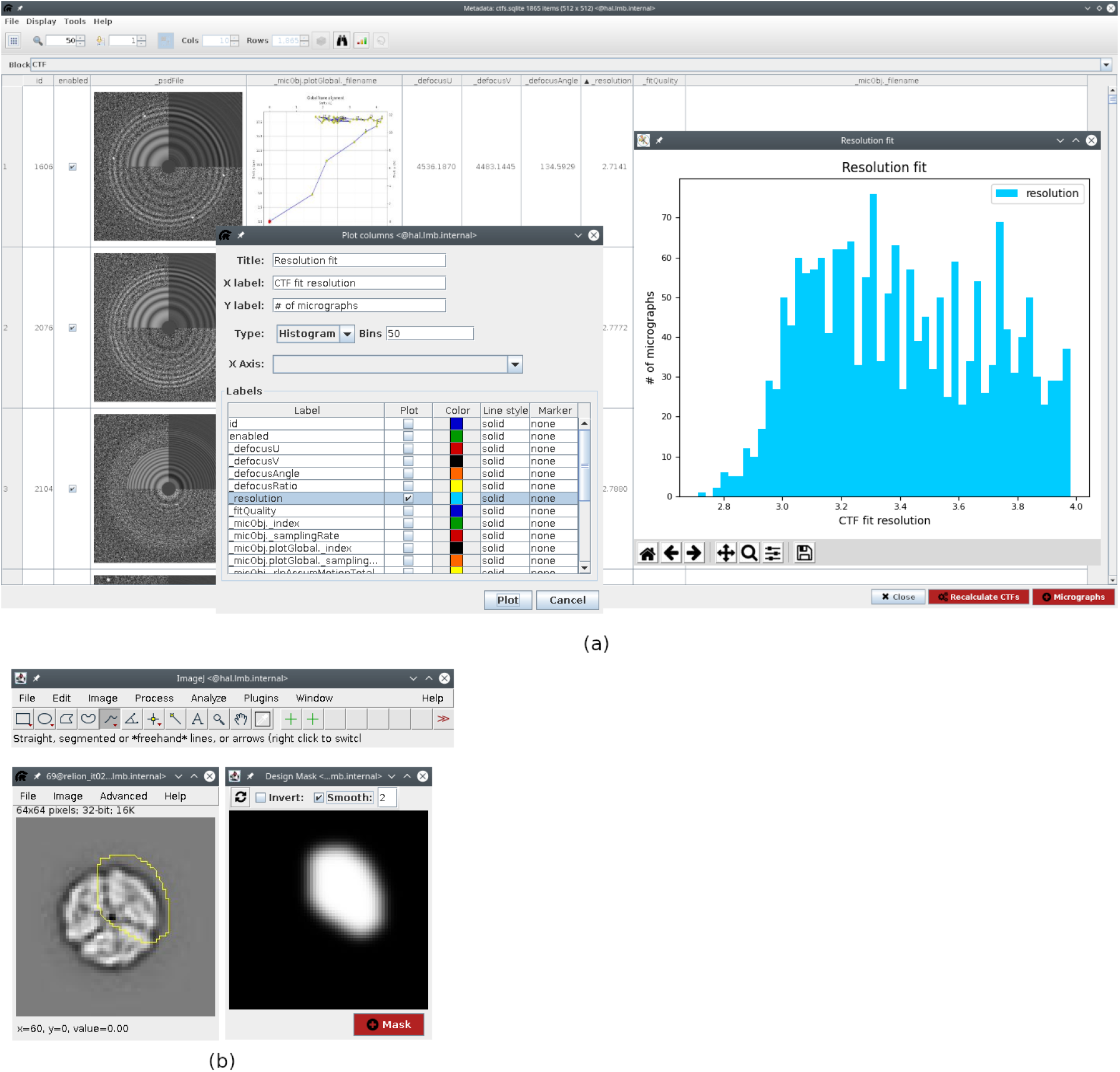
Scipion offers a rich viewing interface for both item sets and individual images. (a) Relion STAR-file metadata can be sorted and plotted in various ways. (b) Individual images can be analyzed and processed with built-in ImageJ tools. Here an interactive wizard for 2D mask design is depicted.

The plugin for Relion also includes custom-designed viewers that not only let users visualize output sets of items from a given protocol, but also create various plots for FSC, SSNR, B-factors, angular distribution and other metrics to quickly follow job progress and iteration results. Protocols for refinement and classification provide the possibility to track changes between iterations, such as orientations, offsets or number of images per class. Other examples include frame motion plots for movie alignment protocols and defocus variation plots for CTF refinement protocols. Combined with the possibilities of creating custom plots from STAR files-derived metadata and displaying volumes in ChimeraX (Pettersen et al., 2020), the plugin for Relion offers a rich interactive interface for both novice and expert users.

## 4. On-the-fly processing

On-the-fly processing (or streaming) is often used for more efficient data collection by providing quick feedback on data quality. It can also considerably shorten the data processing time by overlapping it with microscope acquisition. Scipion provides various options for running workflows in the streaming mode some of which are discussed below.

Firstly, most import protocols have a “Streaming” tab where users can specify a file timeout and a global timeout. The former is used to recognize when the input file size stops growing and is ready to be imported, while the latter is used to stop streaming once no new data is being acquired. Secondly, for several plugins that can run on GPUs Scipion provides a possibility for parallel multi-GPU execution. Such examples are motioncor2 (Zheng et al., 2017) for movie alignment and GCTF (Zhang, 2016) for CTF estimation. By specifying a number of threads equal to the number of GPUs plus one, a user can submit streaming jobs to several GPUs simultaneously. For instance, setting the number of threads to 3 and the GPU IDs to “0, 1” will lead to thread 1 being a master, while thread 2 will run on GPU 0 and thread 3 on GPU 1. Each worker thread will process a single movie in streaming giving a speed boost in processing. More complex combinations are also possible but depend on the underlying protocol.

Multi-GPU execution, however, is different for Relion protocols where a user usually has to specify both MPI processes and threads. We have left the syntax for GPU IDs the same as in the original Relion GUI to avoid confusion. Users can refer to the Relion website documentation for more details.

The protocols for CTF estimation, particle extraction and auto-picking in most Scipion plugins also have a “Streaming” tab where a user can provide a batch size. By default (batch size 1), all input protocol items will be processed one by one which is common for on-the-fly processing pipelines. However, users can specify a larger number of items that will be then processed “in batches”, which sometimes can minimise computational load and speed up processing. Setting this value to 0 is used for non-streaming cases when all input items are processed together.

Altogether, multiple plugins in Scipion support various streaming scenarios for most image preprocessing steps up to 2D classification. Such protocols can be found in the protocol search window of the Scipion project (available by pressing Ctrl+F). Below we show an example of streaming processing on real data.

## 5. Interaction with other software

Besides the ability to import data from various cryo-EM packages provided by Scipion itself, the plugin for Relion includes several protocols that can export results into a self-contained folder, allowing users to continue image processing outside Scipion. Export at the level of coordinates, CTFs (micrographs with CTF information) and particles is possible. We are working towards running all export protocols in streaming so that results could be immediately accessed by other software packages or by Relion itself (outside Scipion).

From the beginning the Scipion framework has always been focused on providing a transparent combination of different software packages. As a result, Relion users are free to incorporate various programs for movie alignment, CTF estimation and particle picking into a standard Relion pipeline. Global movie alignment parameters produced by Motioncor2 (Zheng et al., 2017) or cisTEM - unblur (Grant et al., 2018) can be used as input for Relion’s bayesian polishing. Moreover, the plugin includes several protocols which do not have their own GUI in the RELION interface. Such examples are centering of class averages, movie compression, gain estimation for movie compression, volume symmetrization and several others.

## 6. Image processing

To demonstrate the capabilities of the plugin for Relion we have chosen the EMPIAR-10389 dataset (Righetto et al., 2020). The deposited dataset consists of two sets of movies: one acquired with beam-image shift (multi-shot) and one without (single-shot). The published workflow leads to a 2Å resolution map and comprises a wide range of protocols including the basic steps from pre-processing to 3D refinement along with more complex image processing tasks such as changing the dataset binning, particle re-centering, joining of different datasets, particle polishing and CTF refinement. We think this workflow provides a good example to illustrate Scipion’s capabilities for image processing. Even though based predominantly on Relion, this example shows the advantages of performing these operations within the Scipion framework, rather than using the Relion GUI. To reduce computational costs we have simplified the pipeline by removing several intermediate steps, however the general workflow outline remains the same (fig. 4). To validate the results reproducibility between Relion and Scipion, we have first run this workflow in the Relion 3.1 GUI (outside Scipion) to replicate the steps authors used in their publication to the best of our knowledge. Both procedures are described below.

**Figure 4.**
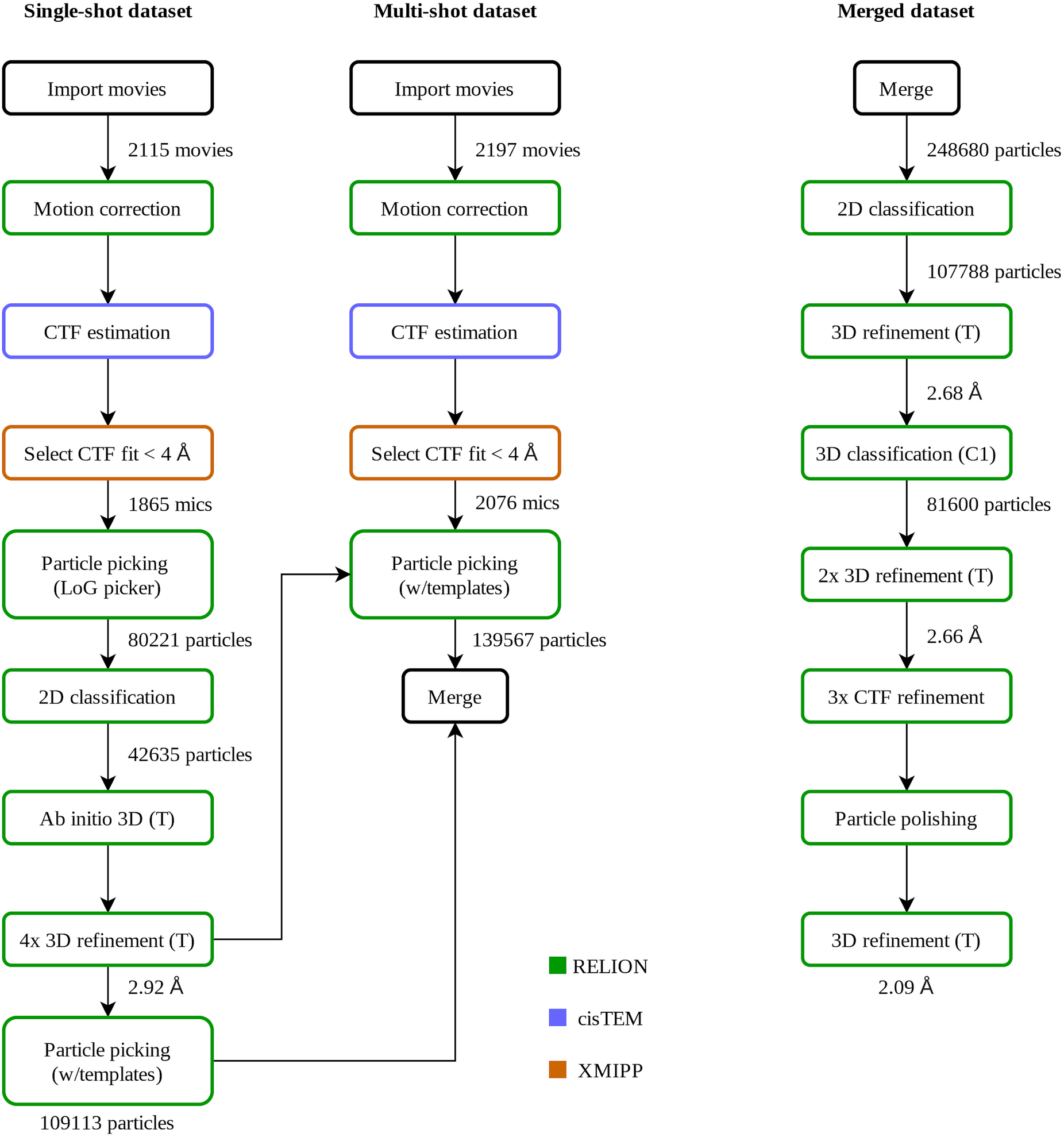
Simplified flow chart of Relion processing within Scipion framework for EMPIAR-10389 dataset. We have first processed the single-shot dataset separately and then used the resulting map for the reference-based particle picking of both single and multi-shot datasets (see text for details). At this point the two datasets were joined and processed further together. For comparison, see Supp. fig. 2 in Righetto et al., 2020.

### 6.1. Processing in Relion outside Scipion

The raw movies were downloaded from EMPIAR (Iudin et al., 2016), split into 4 optic groups based on the movie filename (3 multi-shot groups and 1 single-shot group) and imported separately into the Relion project. Movie STAR files for multi-shot groups were joined together and then both multi and single-shot datasets were processed in the following manner. Movies were motion-corrected with Relion motioncor (no frame grouping, using 5×5 tiles) and amplitude power spectra sums were produced for further CTF estimation with CTFFIND 4.1.14 (Rohou and Grigorieff, 2015). Micrographs with estimated CTF fit lower than 4Å were excluded from further processing. The single-shot dataset was first processed separately.

Initially, the LoG-picker algorithm was used to pick particles, which were then extracted using a 512 pixel box size and binned 8 times to a pixel size of 5.11Å in order to speed-up the downstream processing. Around 80,000 binned particles were classified into 80 2D classes with the “Ignore CTF until first peak” option enabled. Particles from the best 2D classes showing relevant high-resolution features were regrouped (using the “Subset selection” job) and subjected to the stochastic gradient descent algorithm implemented in Relion to generate an *ab initio* 3D map (applying tetrahedral symmetry). The resulting map was used as a reference for the masked 3D refinement that reached the Nyquist limit for the binned data. The particles were then re-extracted with a binning of 4 times using re-centered coordinates by applying shifts from the refinement, and the 3D refinement was repeated. Re-extraction and refinement cycles were repeated until the unbinned box size was used, leading to a 2.84Å reconstruction from ∼ 45,000 particles. In comparison with the publication, we did not perform further refinements, 3D classification or Bayesian polishing for this dataset.

Afterwards we used a reference-based Relion particle picker with the 3D reference map from the last 3D refinement with the data binned twice. We picked approx. 97.000 and 129.000 particles from the single and multi-shot datasets, respectively. The particles were separately extracted, joined and 2D classified (again binned 8 times with a 64 pixel box size to speed-up calculations). Then the particles were extracted again (binned twice) and compared with their counterparts (binned 8 times). Here we had to modify the STAR files manually so they would point to the best particle subset after the 2D classification but with a different binning level. This particle subset was submitted for a 3D refinement that led to a 2.66Å structure from a total of ∼ 118.000 particles. Next, we performed a masked 3D classification (without imposing symmetry) without alignment to further clean the data. Finally, all particles from the best looking and largest 3D class were re-extracted unbinned and refined to a 2.6Å map. We continued the processing with three sequential rounds of CTF refinement to correct for magnification anisotropy, to refine the defocus per particle and astigmatism per micrograph and to correct for high-order aberrations, respectively. This was followed by Bayesian particle polishing using the parameters from the published manuscript. A final round of 3D refinement and post-processing led to a 2.08Å map. This is above the 1.98A reported in the publication. We believe the difference to be caused by the fact that we did not iterate 3D refinements after each CTF refinement step which could have improved the angular accuracy.

### 6.2. Processing in Scipion

After completing the workflow in the Relion GUI, we have switched to the Scipion plugin for Relion. Here all protocols parameters were set consistent with the Relion values used above. Similarly, we have first focused on the single-shot dataset. Gain references and defect files were downloaded from EMPIAR in advance and the preprocessing was performed in streaming mode. As movies were being downloaded the following steps were executed for each movie: import, Relion’s motion correction, CTF estimation with ctffind4 (via the plugin for cisTEM), sorting micrographs by CTF fit (via “ctf consensus” protocol of the plugin for XMIPP), Relion’s LoG picking and particle extraction.

Scipion offers several options for the workflow’s execution: a) importing a predefined workflow file in JSON format into a Scipion project and then running it (done via Scipion GUI or through the command line (https://scipion-em.github.io/docs/docs/facilities/facilities-workflows.html)); b) scheduling protocols to run one after another using the waiting list option; c) creating a workflow interactively: as soon as a single output item of one protocol appears, it can be connected to another protocol as input. Additionally, it is also possible to process items in batches in streaming mode as described earlier. Here we used a combined approach and set up Scipion to run import and motion correction on a one-by-one item basis while CTF estimation, picking and extraction were run in batches on every 20 micrographs. Further processing of the single-shot dataset up to the final 3D refinement with unbinned data was done using the exact same job parameters as for Relion. Rescaling of 3D reference volumes was done with the plugin for XMIPP. Particle re-extraction was performed in a different way compared to Relion. Currently, the particle extraction protocol in the plugin for Relion does not support input particles, so a user must run an “extract coordinates” protocol, which produces a new set of coordinates taking into account particle shifts, in our case from the 3D refinement. This is followed by a normal particle extraction step. Final 3D refinement for the single-shot dataset reached a 2.87Å resolution.

For the multi-shot dataset the preprocessing was done in a standard non-streaming way as joining item sets currently is not possible on-the-fly in Scipion. Similarly to the Relion workflow described above, particles from both single and multi-shot datasets were picked using a 3D reference volume from the previous 3D refinement with the data binned twice. The 8 times binned particles from both datasets were extracted, joined and submitted to 2D classification. Then twice binned particles were re-extracted, joined and cross-referenced with their more binned counterparts using the “pwem - subset” protocol with intersection option. Here no STAR file editing was required to compare particles with different binnings nor to select the best subset after classification. Scipion is able to compare any two item sets as long as they belong to the same “type”, in our case, particles. Afterwards, 3D classification without alignment, followed by re-extraction of unbinned particles from the best class and subsequent 3D refinement was performed resulting in a 2.60Å structure. Before starting CTF refinements we used “relion - assign optics” protocol providing a simple STAR file with two columns, movie name and optics group number, as input to tell Relion that we have 4 different beam tilt classes in our particle set. When using Relion GUI, this step is done at the import stage or by manually editing the STAR files. In Scipion we provide a semi-automated solution where optic groups can be assigned at any step of the workflow. Once all three CTF refinement rounds were completed, we joined the single and multi-shot movies after motion correction for and assigned optic groups again (now to the movies), since particle polishing in Relion also requires movies as input. The final 3D refinement with polished particles reached 2.14Å resolution. We find the final resolution difference between the Relion workflows inside or outside Scipion negligible, which is expected as Scipion wrapper essentially executes the exact same Relion commands provided that input parameters are the same. Minor variations can be attributed to variations in user-based class or particle selections, different 3D refinement starting seeds or small variations in the shifts application during particle re-extraction.

## 7. Conclusions

Here we describe the functionality of the Scipion plugin for Relion software with the focus on single particle image processing for cryo-EM. The plugin provides a set of Python wrappers for the Relion GUI and command-line programs while at the same time offering all capabilities of Scipion’s image and metadata handling via a user-friendly graphical interface. More documentation on the Scipion software can be found at https://scipion-em.github.io/docs/. The plugin code is freely available on Github (https://github.com/scipion-em/scipion-em-relion) and distributed under the GNU general public license (GPL3). For user questions and feedback, please use Github Issues or the Scipion mailing list at https://sourceforge.net/projects/scipion/lists/scipion-users.

## References

1. Bharat, T. A. M., Russo, C. J., Löwe, J., Passmore, L. A., Scheres, S. H. W. (2015). Structure, 23(9), 1743–1753.

2. Danev, R., Yanagisawa, H., Kikkawa, M. (2019). Trends in Biochemical Sciences, 44(10), 837–848.

3. de la Rosa-Trevín, J. M., Otón, J., Marabini, R., Zaldívar, A., Vargas, J., Carazo, J. M., Sorzano, C. O. (2013). J Struct Biol., 184(2), 321–328.

4. de la Rosa-Trevín, J. M., Quintana, A., Del Cano, L., Zaldívar, A., Foche, I., Gutiérrez, J., Gómez-Blanco, J., Burguet-Castell, J., Cuenca-Alba, J., Abrishami, V., Vargas, J., Otón, J., Sharov, G., Vilas, J. L., Navas, J., Conesa, P., Kazemi, M., Marabini, R., Sorzano, C. O., Carazo, J. M. (2016). J Struct Biol., 195(1), 93–99.

5. de la Rosa-Trevín, J. M. and Sharov, G. (2020). DOI: 10.5281/zenodo.4303966.

6. Grant, T., Rohou, A., Grigorieff, N. (2018). eLife, 7, e35383.

7. He, S., Scheres, S. H. W. (2017). J Struct Biol., 198(3), 163–176.

8. Iudin, A., Korir, P. K., Salavert-Torres, J., Kleywegt, G. J., Patwardhan, A. (2016). Nature Methods, 13, 387–388.

9. Pettersen, E. F., Goddard, T. D., Huang, C. C., Meng, E. C., Couch, G. S., Croll, T. I., Morris, J. H., Ferrin. T. E. (2020). Protein Sci, Sep 3.

10. Righetto, R. D., Anton, L., Adaixo, R., Jakob, R. P., Zivanov, J., Mahi, M.-A., Ringler, P., Schwede, T., Maier, T., Stahlberg, H. (2020). Nature Communications, 11, Article number 5101.

11. Rohou, A., Grigorieff N.. (2015). J Struct Biol., 192(2), 216–221.

12. Schneider, C. A., Rasband, W. S., Eleceiri, K. W. (2012). Nature Methods, 9(7), 671–675.

13. Zhang, K. (2016). J Struct Biol. 193(1), 1–12.

14. Zheng, S. Q., Palovcak, E., Armache, J.-P., Verba, K. A., Cheng, Y., Agard, D. A. (2017). Nature Methods, 14, 331–332.

15. Zivanov, J., Nakane, T., Scheres, S. H. W. (2020). IUCrJ, 7(2), 253–267.

16. The Electron Microscopy Data Bank (EMDB) statistics, https://www.ebi.ac.uk/pdbe/emdb/statistics_main.html/

17. Scipion documentation, https://scipion-em.github.io/docs/docs/facilities/facilities-workflows.html

18. Scipion protocol usage ranking, http://scipion.i2pc.es/report_protocols/protocolTable/

