## Supplementary information for "Using Relion software within Scipion framework"

### Differences with the published EMPIAR-10389 workflow

1. Single-shot processing was simplified: we did not perform further refinements (CTF, polishing, classification, etc.) after the first 3D refinement.
2. We have used Relion reference-based picker with the 3D volume as a reference, while in the original manuscript Gautomatch (Zhang, unpublished) was used.
3. We have used the Relion implementation of motioncorr instead of motioncor2. 5x5 patches and no frame grouping were used in our case. Additionally, amplitude power spectra were calculated and used for CTF estimation, while in the manuscript authors used aligned movies for CTF estimation.
4. The deposited EMPIAR datasets do not contain movies for which the CTF resolution fit was worse than 4Å.
5. We used solvent-flattened FSC during all 3D refinements.
6. For the merged dataset we did not perform 3D refinements after each CTF refinement step.
7. During CTF refinement we have opted to refine astigmatism only per micrograph and not per particle.
